# *Easi*-CRISPR: Efficient germline modification with long ssDNA donors

**DOI:** 10.1101/069963

**Authors:** Rolen M. Quadros, Masato Ohtsuka, Donald W Harms, Tomomi Aida, Ronald Redder, Hiromi Miura, Guy P. Richardson, Mark A. Behlke, Sarah A. Zeiner, Ashley M. Jacobi, Lisa D. Urness, Suzanne L. Mansour, Channabasavaiah B. Gurumurthy

**Author notes:** Contributed equally (Rolen M. Quadros, Masato Ohtsuka, Donald W Harms).

## Abstract

CRISPR/Cas9 technology efficiently produces short insertions or deletions (*indels*) and can insert short exogenous sequences at Cas9 cut sites. However, targeting long inserts is still a major technical challenge. To overcome this challenge, we developed *Easi*-CRISPR (*<u>E</u>*fficient *<u>a</u>*dditions with *<u>s</u>*sDNA *<u>i</u>*nserts-CRISPR), a method that uses long, *in vitro*-synthesized, single-stranded DNAs with 50-100 base homology arms as repair templates. We demonstrate that *Easi*-CRISPR can generate knock-in and floxed alleles in mice with an efficiency at many loci as high as 100%. The simple design requirements for donor DNAs and the reproducibly high-efficiency of *Easi*-CRISPR enables rapid development of many types of commonly used animal and cell models.

## Introduction

The CRISPR/Cas9 system is used in many fields of research to create genome-modified cells and organisms. It is routinely used to create short insertions or deletions *(indels)* via non-homologous end-joining (NHEJ) and also to insert sequence content from short singlse-stranded oligodeoxynucleotides (ssODNs) into genes of interest via homology directed repair (HDR). The ssODN repair templates are typically about 100-200 bases long, consisting of a few bases of altered sequence (e.g., point mutations, recombinase recognition sequences, short deletions or insertions of a few bases) flanked by homology arms of about 40-80 bases (1,2). Targeted insertion of longer new sequences (>100 bases) typically employs cloned double-stranded DNA (dsDNA) as the repair template because ssODNs longer than 200 bases cannot be obtained commercially. However, in comparison with ssODN donors, the efficiency of dsDNA template insertion is often poor (3,4). Furthermore, dsDNA templates require homology arms of at least 0.5 kb (4,5). The technical constraints of designing and building such custom targeting constructs for each project add limitations to this approach.

Some strategies for increasing the targeting efficiency of donor DNAs include inhibition of NHEJ or enhancing HDR through chemical treatments (6–10). Such methods are based, however, on perturbation of fundamental DNA repair processes and may be toxic. Non-toxic approaches include the use of circular dsDNA donors with built-in artificial guide sequences that are linearized inside the cell/embryo (11–13). The linearized donor DNA is then inserted at the genomic Cas9 cut site by cellular ligases. These designs include either micro-homology ends between the cut ends of the genomic DNA and donor DNA, or ssODNs that bind to the two cut ends so that a precise fusion occurs between the donor and genomic DNAs. While these strategies offer better alternatives to those that perturb DNA repair, they too have limitations, including the need to design special donor plasmids to suit each target site.

The two most common types of knock-in animals needed for biomedical research and molecular genetic studies are those carrying: i) targeted insertions of commonly used sequences encoding recombinases such as Cre and Flp; transcriptional regulators such as rtTA and tTA; and reporters such as EGFP and tdTomato; and importantly: ii) Conditional Knock-Out (CKO) alleles in which two *loxP* sites flank one or more critical exons (i.e. “floxed” alleles). The mouse research community still depends on lengthy ES cell-based approaches to develop such models because simple CRISPR-based strategies are not available for efficient insertion of longer foreign DNAs.

To develop a simple method suitable for generating many common types of knock-in animal models, we tested a long ssDNA repair template strategy in mice. The rationale was as follows: short ssODN templates (up to 200 bases) are quite efficient in directing insertions at Cas9 cut sites and the same repair-mechanism might be exploited for delivering larger cargo if the length of the ssDNA could be extended. We demonstrated recently that ~500 base ssDNAs containing ~400 bases of synthesized microRNA sequence flanked by ~55 base homology arms directed efficient targeting of the mouse genome (10-83% efficiency when analyzed during embryonic stages) (14). In the present study, we tested the suitability of this method for i) inserting 1-1.5 kb protein coding cassettes into different genes, and ii) generating floxed alleles. The method reported here, termed *Easi*-CRISPR (*<u>E</u>*fficient *<u>a</u>*dditions with *<u>s</u>*sDNA *<u>i</u>*nserts-CRISPR; pronounced *Easy*-CRISPR) can be readily adapted to generate even longer insertions at Cas9 cut sites.

We developed four knock-in mouse strains using *Easi*-CRISPR. Three models consisted of an ~1.4 kb cassette encoding P2A-FlpO recombinase inserted immediately before the stop codons of *Fgf8, Slc26a5* and *Mafb.* The fourth model consisted of ~1 kb reverse tetracycline transactivator (rtTA)-polyA cassette inserted in-frame immediately after the initiation codon of *Otoa.* In each case, the cassettes contained flanking homology arms to the target loci that ranged between 72 and 105 bases. Long ssDNA donor molecules were synthesized using the *IvTRT* (*in vitro* Transcription and Reverse Transcription) method described previously (14), or obtained from a commercial source (single-stranded gBlock^®^ gene fragments, Integrated DNA Technologies). Schematics of the ssDNA cassettes are shown in Figure 1a, Supplementary Figure 1a, 2a and 3a and their full sequences are shown in Supplementary Figures 4 to 7.

**Figure 1.**
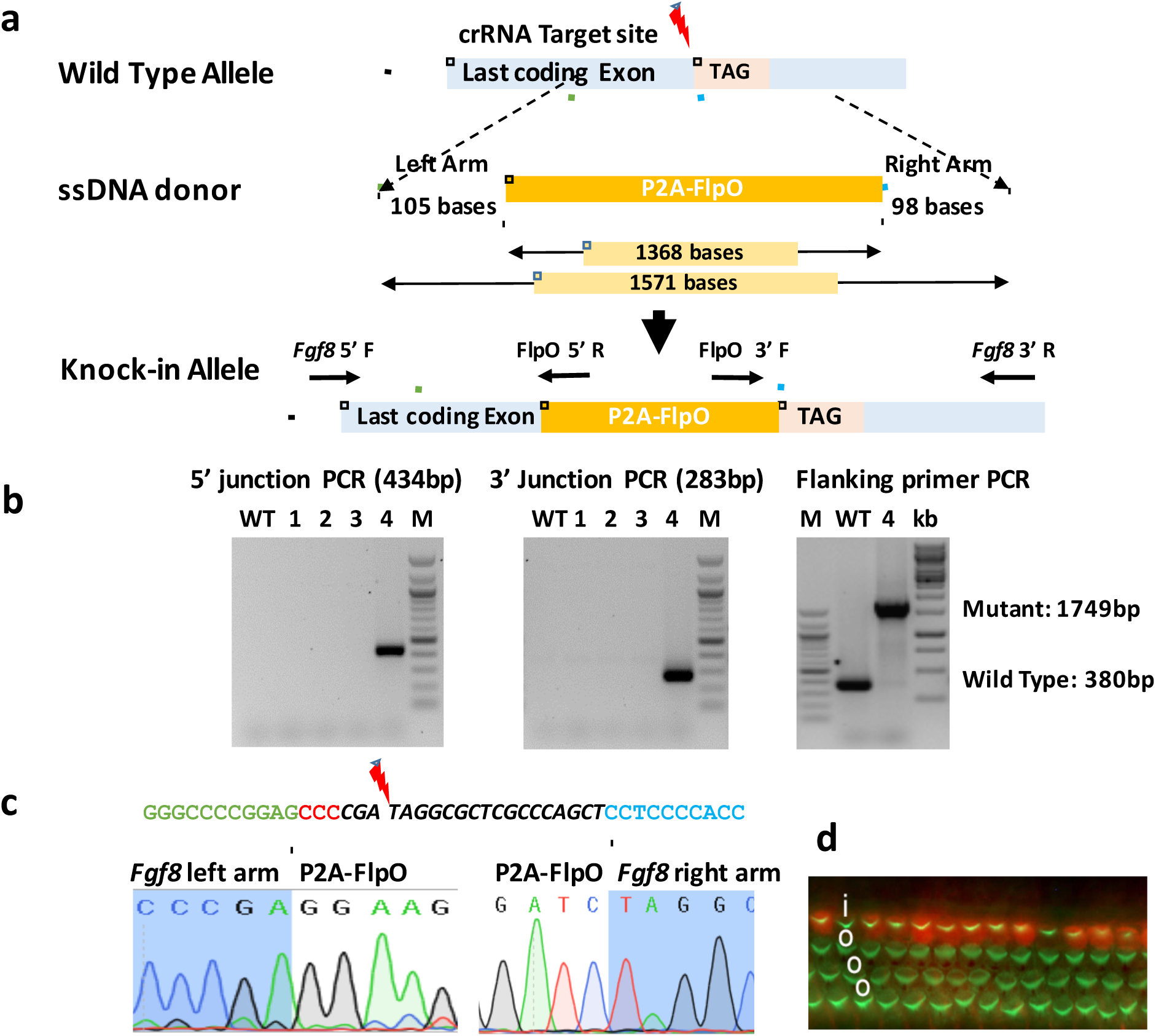
Fusing P2A-FlpO cassette to the 3′ end of *Fgf8* gene using *Easi*-CRISPR.

**(a)** Schematic showing *Fgf8* locus, ssDNA donor and resulting targeted insertion allele. **(b)** Genotyping of GO animals. Schematics of primer locations for 5′ and 3′ junction PCRs are shown along with the expected amplicon sizes. Founder #4 has a correctly targeted P2A-FlpO insertion, as indicated by the presence and size of both 5′ and 3′ junction amplicons. The gel on the right shows that PCR amplification of this founder’s DNA with primers flanking the *Fgf8* insertion site produced only the mutant amplicon, indicating that it is a biallelic insertion. WT: wild type, M: 100 bp marker, kb: 1 kb Marker. **(c)** Sequencing of 5′ and 3′ junctions in founder #4. The guide RNA sequences (italics), along with the cut sites, PAM sequences (in red) and a few bases of flanking sequences (above) and sequence chromatograms showing correctly targeted 5′ and 3′ junctions (below). **(d)** FGF8-P2A-FLPO activates a FLP-dependent tdTomato reporter in inner hair cells. Surface preparation of the cochlear epithelium isolated from a P1-P2 *Fgf8^p2A-Flpo/+^;Rosa26^RC::RFLG/+^* pup was stained with Alexa488-phalloidin (green). Native tdTomato fluorescence (red) is evident in most inner hair cells (i), but not in outer hair cells (o).

The founder G0 pups were generated following standard CRISPR/Cas9 mouse genome engineering protocols (15) with some modifications described in the online methods. Details of the microinjections and the results are shown in Supplementary Table 1. Genotyping of G0 pups using 5′ and 3′ primers (Supplementary Table 2) indicated that the overall insertion efficiency at all four loci was 28% (7 out of 25 pups contained the respective insertion cassettes; Figure 1b, Supplementary Figures 1b, 2b and 3b). The individual insertion efficiencies, for different genes, ranged from 25% to 33% (Supplementary Table 1). Notably, one founder contained bi-allelic insertions of the knock-in cassette (Figure 1b). The fidelity of the insertions, including the junctions, was confirmed by sequencing (Figure 1c and Supplementary Figure 1c, 2c and 3c). All targeted animals contained precisely targeted insertions, except for one *Slc26a5^P2A-FlpO^* founder in which genomic DNA sequence duplications were present at the 3′ end (Supplementary Figure 1b). Two of the founders tested to date produced heterozygous F1 offspring (Supplementary Figure 8). Furthermore, F1 offspring of the *Fgf8-P2A-FlpO* founder #4 showed the expected expression of FLPO in cochlear inner hair cells (Figure 1d). Taken together, these results indicate that *Easi*-CRISPR can insert sequences of ~1 kb or longer at high efficiency and that the technique is reproducible at multiple genomic loci.

CKO mice are among the most highly used genetically engineered models, but unfortunately there are no compelling strategies for generating such models using the CRISPR/Cas9 system. In order to test whether *Easi*-CRISPR can generate floxed alleles, we chose three candidate genes. To achieve this goal, we designed two guide RNAs to cut and replace the selected genomic exons with a floxed exon cassette. The details of the target exons, lengths of the ssDNA repair templates, and genotyping strategies are given in Figure 2a, supplementary Figures 9a and 10a. Their full sequences are shown in Supplementary Figures 11–13 and the microinjection details are given in Supplementary Table 1. We injected Cas9 protein and separated crRNA and tracr RNAs as ribonucleoprotein (RNP) complexs for each of three floxed exon cassettes (from the *Col12a1, Paf1* and *Ppp2r2a* genes) (17). Genotyping of the 9 founder pups born (three for each gene) showed that all pups contained at least one floxed allele, a surprising 100% efficiency. Two of these pups also contained biallelic insertions of the cassettes (Figure 2b; sample #3 and Supplementary Figures 10b; **sample #3 and** Supplementary Table 1). The fidelity of the inserts and the correct fusions were confirmed by sequencing (Supplementary Figures 11–13). We observed that one of the pups contained only one *loxP* site, suggesting that partial insertion of the ssDNA cassette can also occur. A similar targeting strategy was designed to flox an exon at a fourth locus but in this case Cas9 mRNA and a standard sgRNA were included in the injection mix. This resulted in some founder pups containing only one of the two loxP sites, indicating partial cassette insertions (data not shown). Notably, all of the floxed alleles that were generated at 100% efficiency were obtained RNP complex microinjection (17),suggesting that the high efficiency of full-length cassette insertion is attributable to RNP injections. However, more experiments involving side-by-side comparisons at the same set of loci, using RNP injections vs Cas9 mRNA and sgRNA injections would be necessary to reach a clear conclusion.

**Figure 2:**
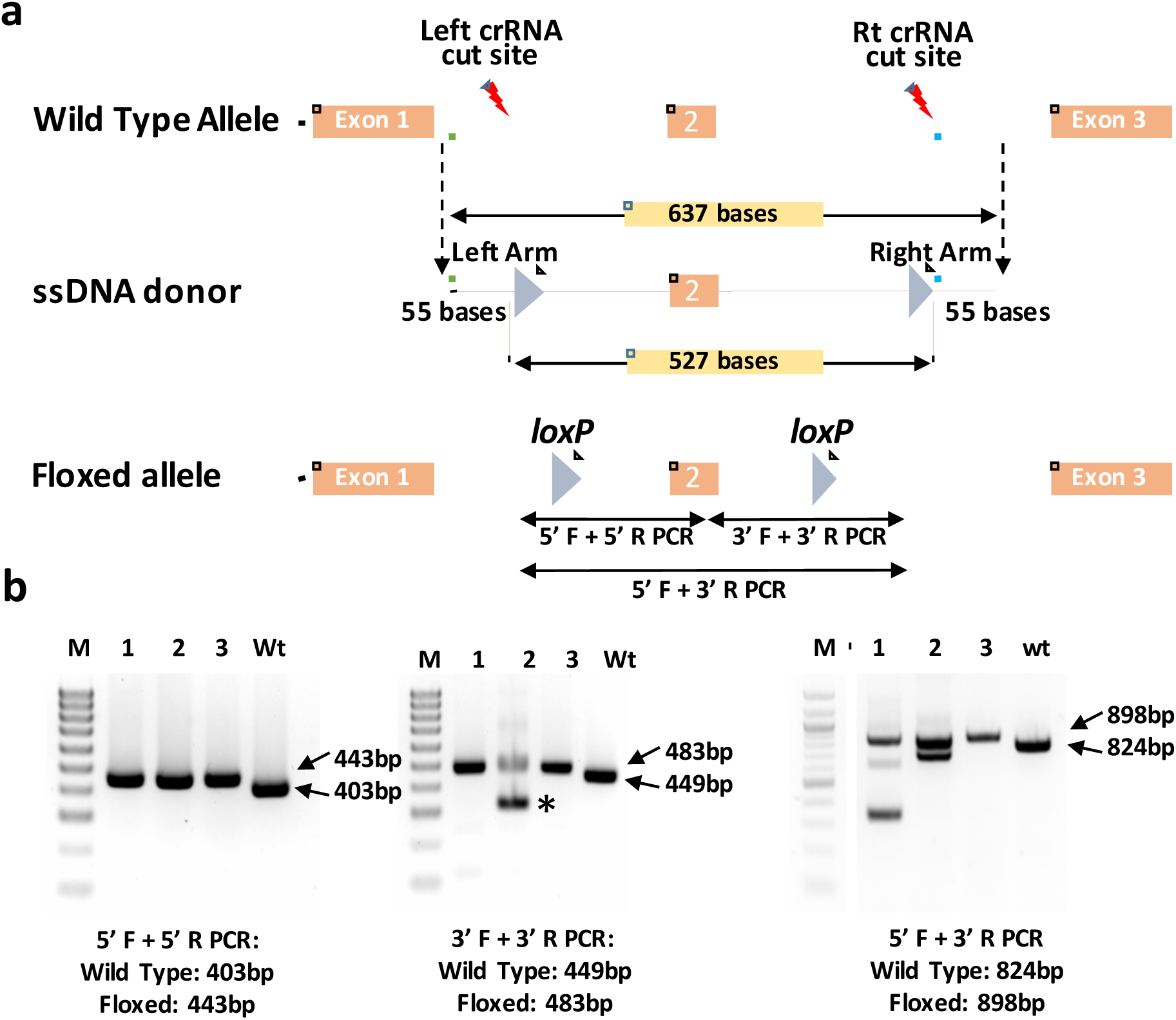
Generation of floxed *Col12a1* allele using *Easi*-CRISPR. **(a)** Schematic showing the wild type allele, ssDNA donor and targeted allele for *Col12a1* locus. The length of the ssDNA, homology arms and the distance between the two *loxP* sites are shown. **(b)** Genotyping of F0 pups for detection of 5′ and 3′ *loxP* insertions. The expected sizes of the PCR amplicons (wild type or floxed) for are indicated. **(c)** *Col12a1* founder #3 was biallelic for both the 5′ and 3′ *loxP* sites, whereas the other two were heterozygous for both 5′ and 3′ *loxP* sites and they carried deletions in their second allele (shown by the asterisk). The gel on the right shows PCR bands from the external primers to both *loxP* sites.

Currently, there are two main strategies for generating floxed alleles; insertion of two *loxPs* in the same allele using two separate ssODNs and two separate sgRNAs (16), or through homologous recombination of long donor dsDNA with at least 0.5 kb of homology arms (4). Both approaches are highly challenging and are not efficient: while the published reports demonstrate about 5 to 16% efficiency (4,16) many labs are still struggling to apply these strategies for other loci. In addition, some of these methods also require specially designed targeting vectors. In contrast, our *Easi*-CRISPR approach i) does not require specially designed targeting vectors, and ii) achieves highly efficient insertion of floxed donor cassettes (up to 100%), especially when RNP-based delivery of CRISPR components is used. Although higher efficiency of targeting when using RNPs with separated crRNA and tracrRNAs instead of a sgRNA has been reported previously (17), to our knowledge, such RNP strategies have not been used in combination with ssDNA repair templates so far. Therefore, we suggest that *Easi*-CRISPR with RNP injections is by far the simplest and most efficient way to create floxed alleles.

Many types of knock-in animal models are routinely used in biomedical research. For example, several thousands of knock-in mice were designed to express commonly used protein-coding sequences, the majority of which are about 1 to 1.5 kb long. Even though the CRISPR/Cas9 system has made several paradigm shifts in animal transgenesis (18), the research community still depends on lengthy ES cell-based approaches to develop knock-in and floxed animal models because there are no simple CRISPR/Cas9-based strategies available for efficient insertion of longer sequences directly in zygotes. Given the simple design requirements for the donor DNA (addition of 50 to 100 base homology arms to a cassette) and the high efficiency of *Easi*-CRISPR (25 to 33% efficiency for insertion of a protein coding cassette at four different genomic loci, and 7 to 100% efficiency for generating floxed alleles at three different loci), this method provides a simple and efficient means of developing many of the commonly used transgenic mouse models and should be readily adaptable to other species or cells in culture.

The *Easi*-CRISPR method could also offer an appealing strategy for inserting multiple point mutations situated in close proximity in the genome as, for example, to test gene regulatory elements. Lastly, whilst the mechanism by which ssDNA molecules are used as templates in DNA repair processes is not well understood, ssDNA-mediated repair mechanisms and the cellular proteins required for such repair processes (19, 20) are emerging. In this context, *Easi*-CRISPR can serve as a tool to interrogate molecular mechanisms of DNA repair by ssDNA donors.

## ACKNOWLEDGMENTS

This work was supported in part by an Institutional Development Award (PI: Shelley Smith) P20GM103471 (to C.B.G, R.M.Q and D.W.H), by NIH R21DC014779 (to S.L.M) and by KAKENHI (26830131, 16K07085, Comprehensive Brain Science Network and Adaptive Circuit Shift) from JSPS and MEXT, grants from Nakatani Foundation, Takeda Science Foundation, MRI, and CNSI/NINS (BS281001) to T.A, and the Wellcome Trust (Grant 087377) to G.P.R. Funding support for generation of floxed alleles for *Ppp2r2a* and *Paf1* genes was provided through NIH R01 CA 138791 (PI: Surinder Batra), NIHK22 CA175260 (PI: Ponnusamy Moorthy) grants, respectively. We thank Harumi Ishikubo, Takako Usami and Yayoi Izu and the genome editing facility at laboratory of recombinant animals, MRI, TMDU, and Y. Wada (FASMAC) for providing technical assistance and materials.

## AUTHOR CONTRIBUTIONS

C.B.G, G.P.R, M.O, and S.M. conceived this study, C.B.G, R.M.Q, G.P.R, M.O, H.M, T.A, M.A.B, and S.L.M. designed the experiments, C.B.G, R.M.Q, D.W.H, T.A, S.A.Z, R.R, L.D.U and A.M.J performed experiments. C.B.G, R.M.Q, G.P.R, and S.L.M. wrote the manuscript with input from all authors.

## Online Materials and Methods

### CRISPR reagents

CRISPR guide RNAs were designed using CRISPR.mit.edu and were used as single guide RNAs (sgRNAs) for *Otoa* or as annealed 2-part synthetic crRNA and tracrRNA molecules for *Fgf8, Slc26a5, Mafb, Paf1, Ppp2r2a* (Alt-R™ CRISPR guide RNAs, Integrated DNA Technologies). The sgRNA was transcribed from a template generated by annealing two primers using the HiScribe™ T7 Quick High Yield RNA Synthesis Kit (New England Biolabs (NEB)) following the manufacturer’s instructions. Cas9 mRNA was prepared using the pBGK plasmid as described in Harms et al., 2014. The plasmid was linearized with XbaI, gel purified and used as the template for *in vitro* transcription using the mMESSAGE mMACHINE T7 ULTRA kit (Ambion: AM 1345). Cas9 protein was obtained from IDT (Alt-R™ S.p. Cas9 Nuclease 3NLS). Single-stranded DNA donors were prepared using the *IvTRT* method described previously (14) or obtained from Integrated DNA Technologies. The floxed *Col12a1* mice were generated using chemically synthesized crRNAs and tracrRNA (Genome Craft Type CT, FASMAC), and Cas9 protein (NEB).

### Microinjection of one-cell embryos

All animal experiments performed were approved by the institutional IACUC protocols. C57BL/6 mice at 3–4 weeks of age (Charles River Laboratories) were superovulated by intraperitoneal injection of 5 IU pregnant mare serum gonadotropin, followed 48 hours later by injection of 5 IU human chorionic gonadotropin (both hormones from National Hormone & Peptide Program, NIDDK). Mouse zygotes were obtained by mating C57BL/6 stud males with superovulated C57BL/6 females. One-cell stage fertilized mouse embryos were injected with 20 ng/µl Cas9 protein (or 10 ng/µl of Cas9 mRNA; for the *Otoa* locus), 20 ng/µl of annealed crRNA and tracrRNA (or 10 ng/µl of each sgRNA; for the *Otoa* locus) and 5-10 ng/µl of ssDNA. Both cytoplasmic and pronuclear injections were performed. The surviving embryos were surgically implanted into pseudo-pregnant CD-1 females.

### Mouse genomic DNA extraction, genotyping and sequencing

Mouse genomic DNA was extracted from toe samples using the Qiagen Gentra Puregene Tissue Kit. Primers were designed to amplify the correctly targeted junctions. Genomic DNA was subjected to flanking primer PCR and internal (donor oligo-specific) and external primer PCR. PCR reactions were performed using the Promega Hot Start green mix. The amplicons were separated on a 1% agarose gel. The gel-purified amplicons were subjected to sequencing using one of the PCR primers.

### FLPo activity assay

A homozygous FLP reporter, B6.Cg-*Gt(ROSA)26Sor^tm1.3(CAG-tdTomato,-EGFP)Pjen^/J* (JAX Stock #026932) (Plumer et al., 2015), was crossed with the *Fgf8*-P2A-FlpO founder #4. Whole cochleae were dissected from P1-P2 pups, cut along Reissner’s membrane to expose the surface of the sensory epithelium and fixed overnight at 4°C in 4% paraformaldehyde made in PBS. The cochleae were stained with Alexa488-phalloidin (Invitrogen) diluted 1:1500 in PBS containing 0.01% Triton X-100 for 15 minutes and then mounted in Fluormount G (SouthernBiotech) on microscope slides. Cochleae were imaged on an Axioskop (Zeiss) with epifluorescent illumination and photographed with an Infinity 3-6UR (Lumenera) digital camera. The green and red channels were overlaid using Photoshop CS6 (Adobe).

---

**Supplementary Table 1:**
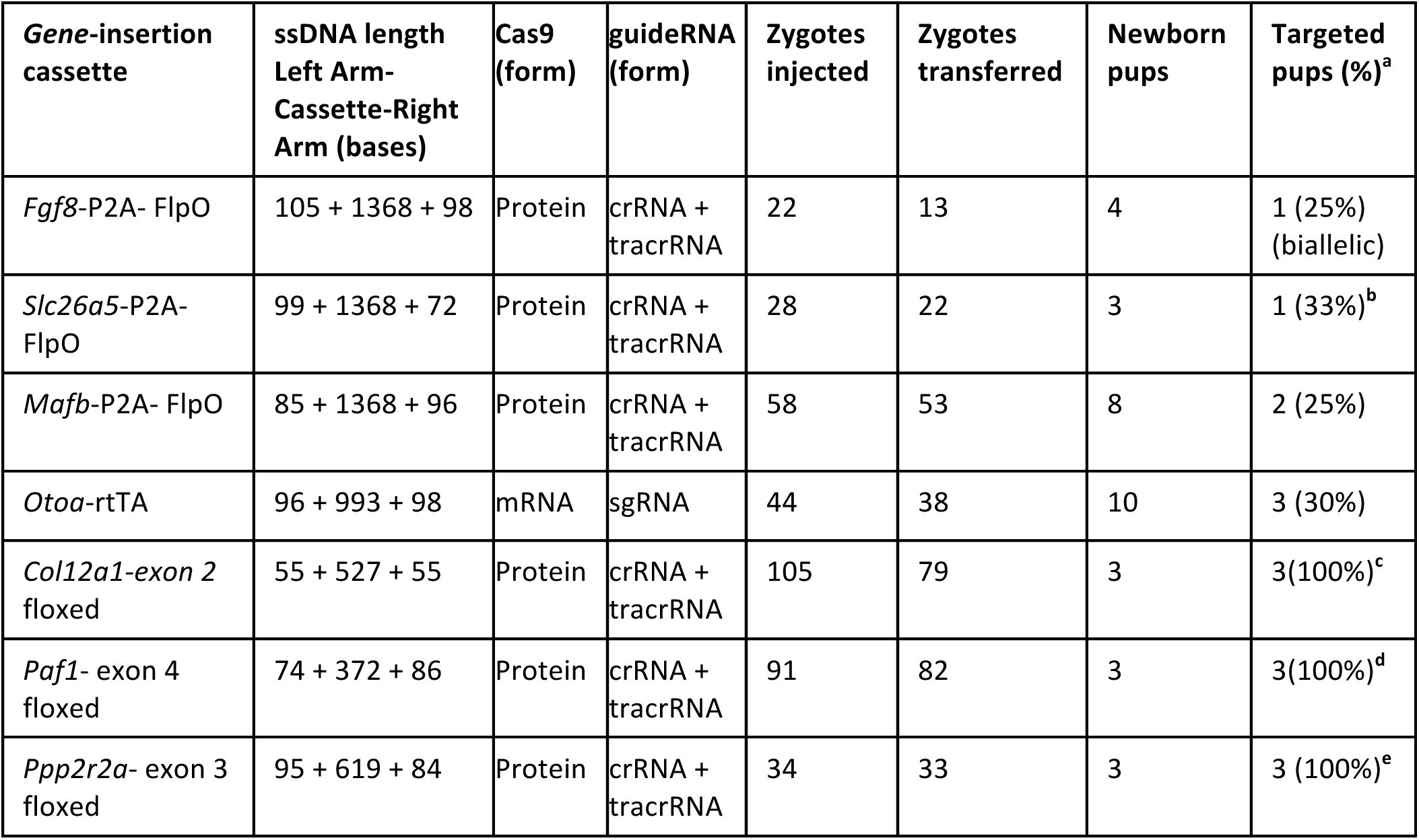
microinjection data:

a. The alleles that did not contain the inserts were not analyzed for presence of *indels* because genotyping assays were designed to identify mainly the targeted-insertion alleles (knock-in or floxed). However, noticeable sized additions or deletions were observed for some samples in these genotyping assays (e.g., additions in Slc26a5 founder #1; Supplementary figure 1b; **lane 1,** and deletions in the non-targeted alleles in floxing experiments; figure 1b and supplementary figures 9b and 10b)
b. Pup #1 had additional sequences at the 3′ junction (sequence not characterized completely) and pup #1 had a precise insertion at both junctions (see Supplementary Figure 1c)
c. The pups #1 and #2 were heterozygous for both 5′ and 3′ *loxP* sites and they carried deletions in their second allele and the pup #3 was biallelic for both the 5′ and 3′ *loxP* sites (see Figure 2b).
d. Pup #1 was heterozygous for both 5′ and 3′ *loxP* sites and the second allele had a deletion; pup #2 was heterozygous for 5′ *loxP* and homozygous for 3′ *loxP* and the pup #3 was heterozygous for both *loxP*s (see Supplementary Figure 9c).
e. Pup # 1 and 2 are heterozygous for both *loxPs* with deletions in the second allele and the pup #3 is biallelic (see Supplementary Figure 10c).

**Supplementary Table 2.**
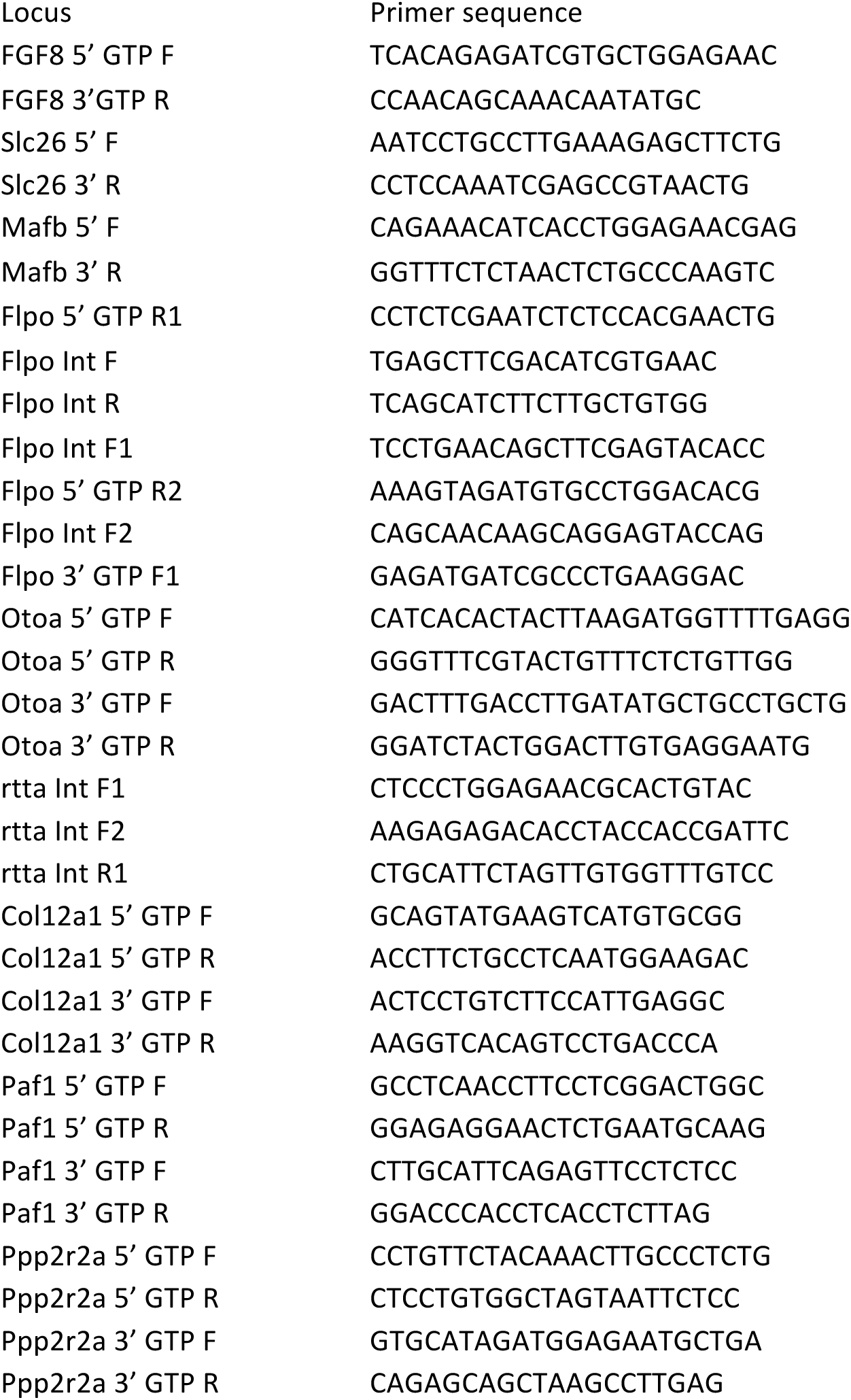
Primer sequences used in this study.

**Supplementary Figure 1:**
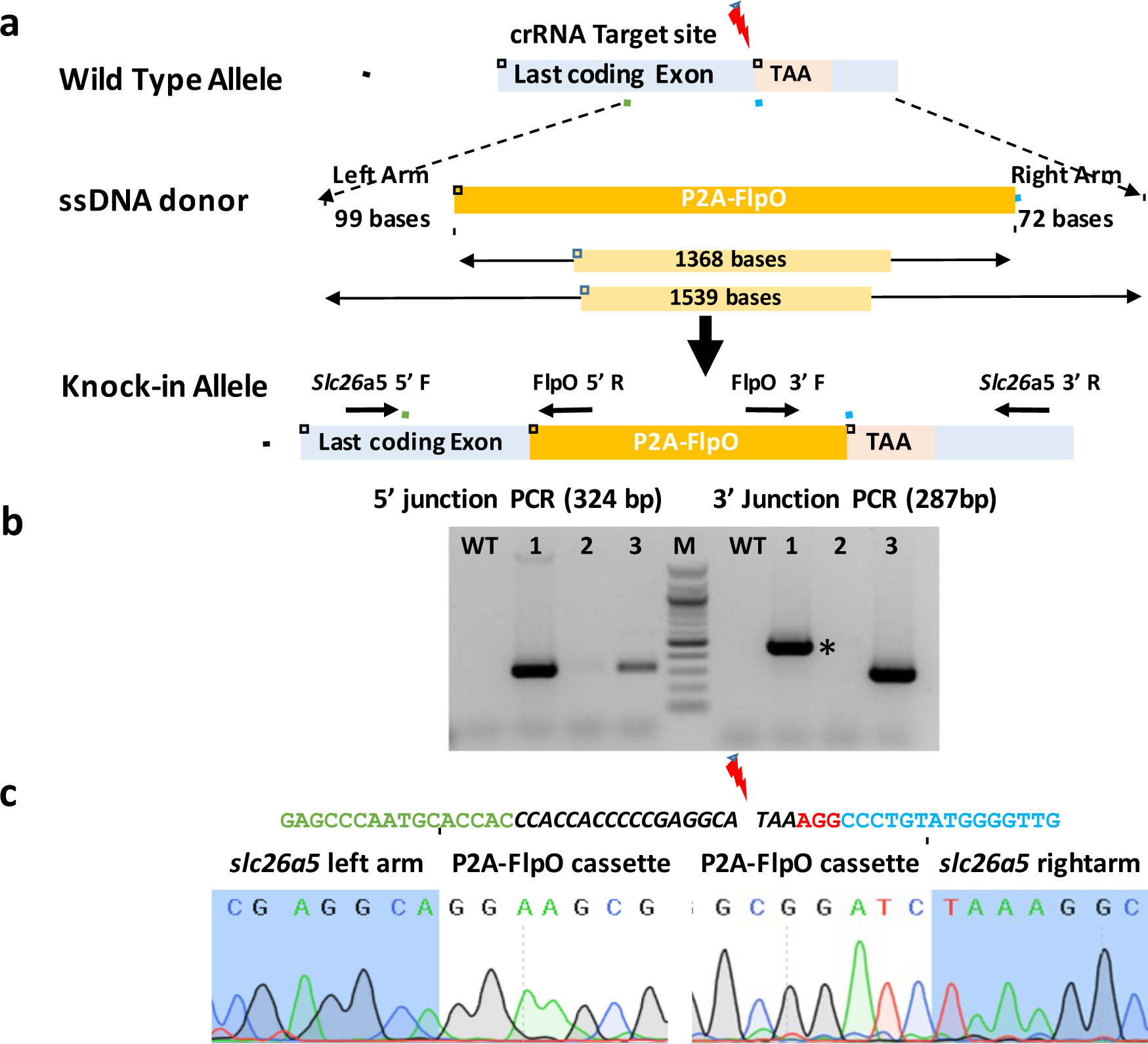
Fusing P2A-FlpO cassette to the 3′ end of *Slc26a5* gene using *Easi*-CRISPR. **(a)** Schematic showing *Slc26a5* ssDNA donor template and targeted knock-in alleles. The lengths of ssDNA, homology arms and the P2A-FlpO cassettes are indicated. **(b)** Genotyping G0 animals for detection of cassette insertion. Schematic of primer locations for 5′ and 3′ junction PCRs is shown along with the expected amplicon sizes. Founders #1 and #3 are positive for the P2A-Flpo cassette insertion using both 5′ and 3′ PCRs. Note that the 3′ junction PCR for founder #1 is bigger than the expected size (shown by an asterisk) suggesting that it contains additional sequences (the exact sequence of the insertion was not determined). **(c)** The guide RNA sequences (italics), along with the cut sites, PAM sequences (in red) and a few bases of flanking sequences are shown above. Sequencing of founder #1 showing correct insertion of the cassette at 5′ and 3′ junctions.

**Supplementary Figure 2:**
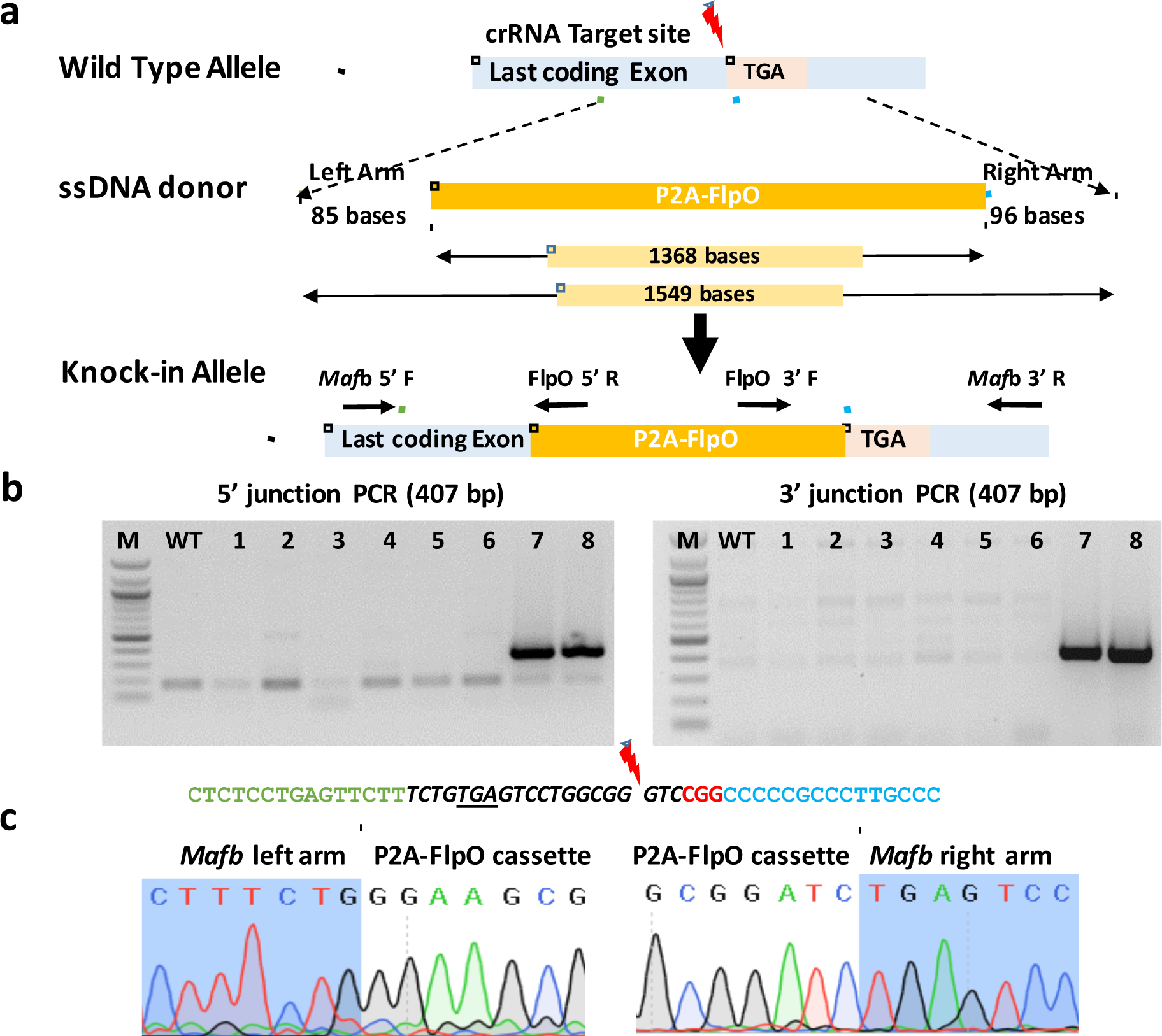
Fusing P2A-FlpO cassette to the 3′ end of *MafB* gene using *Easi*-CRISPR. **(a)** Schematic showing *Mafb,* ssDNA donor template and targeted knock-in alleles. The lengths of ssDNA, homology arms and the P2A-FlpO cassettes are indicated. **(b)** Genotyping G0 animals for detection of cassette insertion. Schematic of primer locations for 5′ and 3′ junction PCRs is shown along with the expected amplicon sizes. Founders #7 and #8 are positive for the Flpo cassette insertion by both 5′ and 3′ junctions PCR WT: wild type, M: 100 bp marker. **(c)** Sequencing founder #7. The guide RNA sequences (italics), along with the cut sites, PAM sequences (in red) and a few bases of flanking sequences are shown above. Note that guide cut site for *Mafb* is 13 bases downstream of the desired stop codon TGA (underlined). Sequence chromatograms showing cassettes insertion junctions.

**Supplementary Figure 3:**
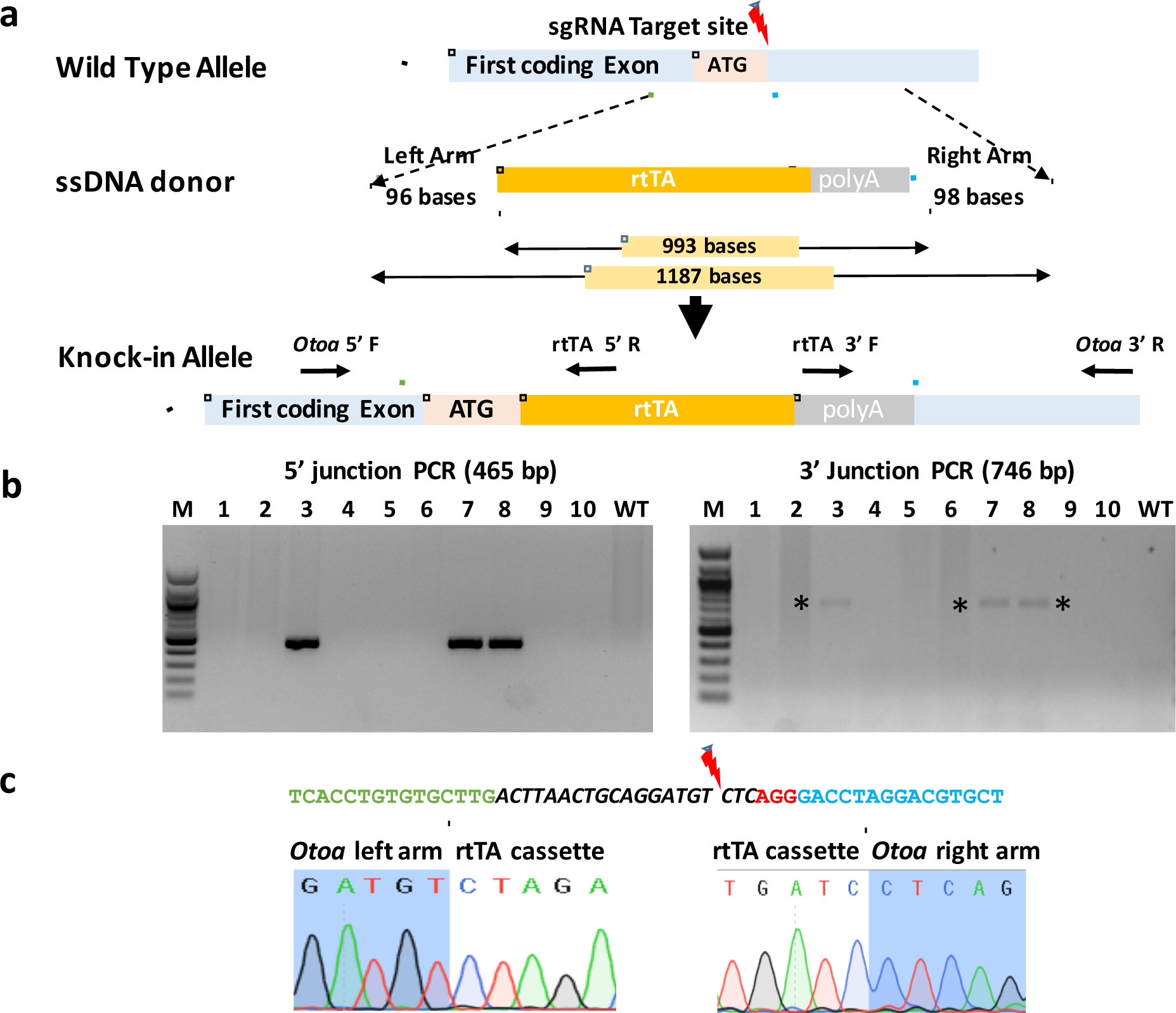
Insertion of rtTA-polyA cassette after the initiation codon of *Otoa.* **(a)** Schematic showing the *Otoa* locus, ssDNA donor and resulting targeted insertion alleles. **(b)** Genotyping of G0 animals. Schematics of primer locations for 5′ and 3′ junction PCRs are shown along with the expected amplicon sizes. Samples from *Otoa* G0 pups 3, 7 and 8 have a correctly targeted rtTA insertion as seen by both the 5′ and 3′ junction PCRs. Note that the amplicon for the 3′ PCR is weak (indicated by asterisks). **(c)** Sequencing of 5′ and 3′ junctions in *Otoa* founder #3 and *Fgf8* founder #4. The guide RNA sequences (italics), along with the cut sites, PAM sequences (in red) and a few bases of flanking sequences (above) and sequence chromatograms showing correctly targeted 5’ and 3’ junctions (below).

**Supplementary Figure 4.**
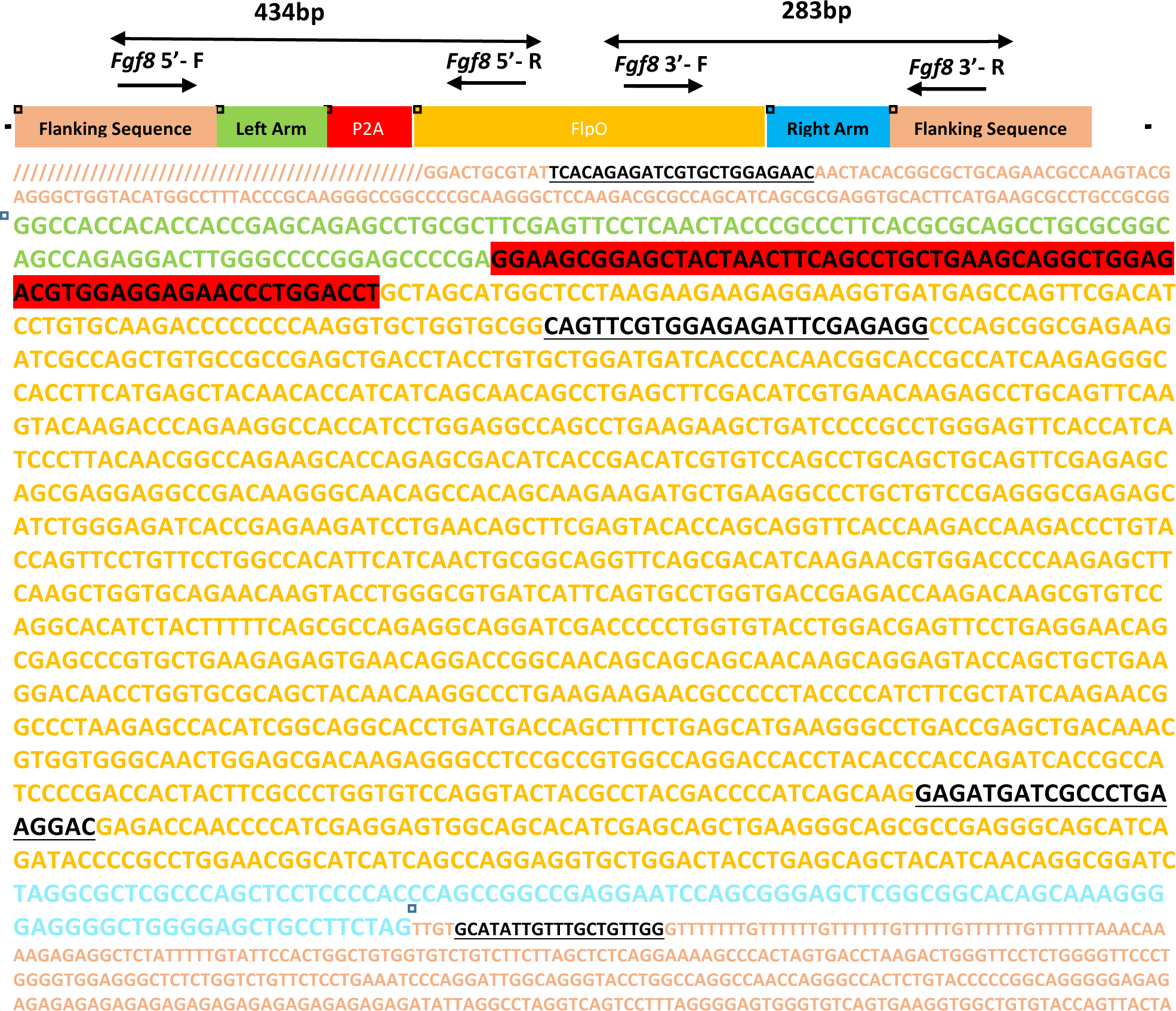
(a) Schematic and sequence of the *Fgf8*-P2A-FlpO-ssDNA. Various sequence elements are color coded. The primers used for 5′ and 3′ junction PCR and the amplicon sizes are shown above the schematic. The full sequence of the ssDNA cassette (including the homology arms) is boxed. Underlined black sequences are primer sequences.

**Supplementary Figure 5.**
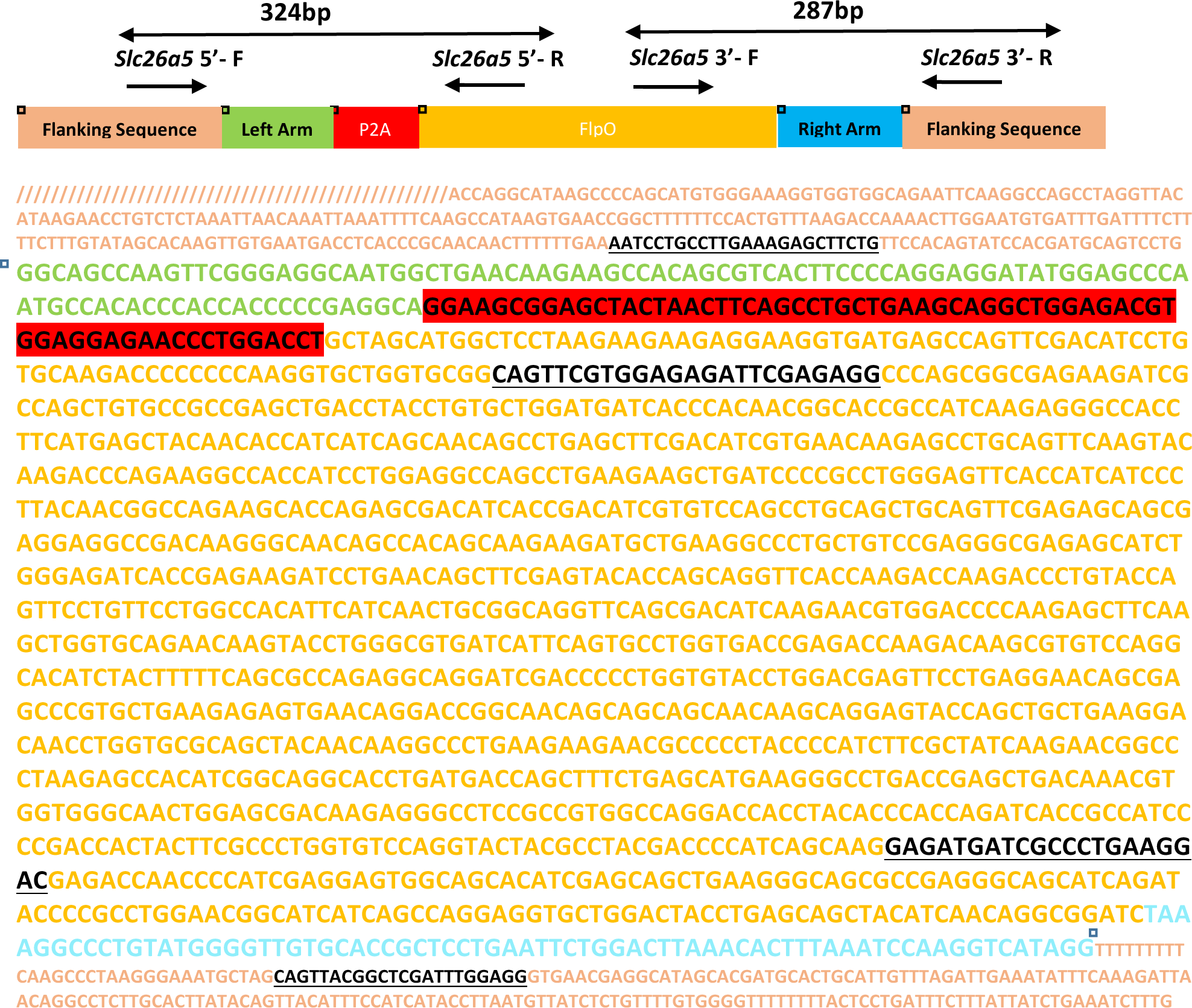
(a) Schematic and sequence of the *Slc26a5*-FlpO-ssDNA. Various sequence elements are color coded. The primers used for 5′ and 3′ junction PCR and the amplicon sizes are shown above the schematic. The full sequence of the ssDNA cassette (including the homology arms) is boxed. Underlined black sequences are primer sequences.

**Supplementary Figure 6.**
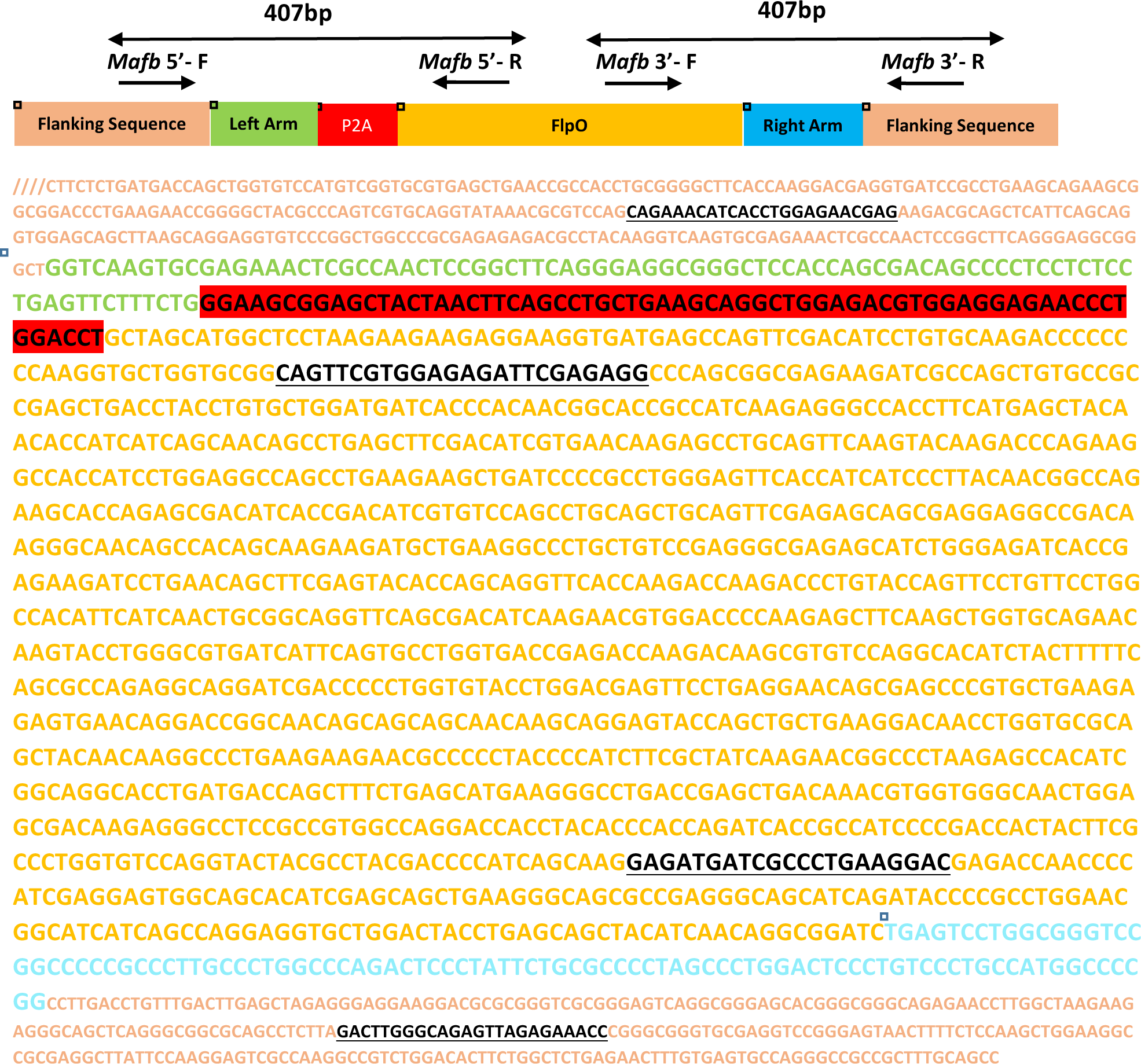
(a) Schematic and the sequence of the *Mafb*-FlpO-ssDNA. Various sequence elements are color coded. The primers used for 5′ and 3′ junction PCR and the amplicon sizes are shown above the schematic. The full sequence of the ssDNA cassette (including the homology arms) is boxed. Underlined black sequences are primer sequences.

**Supplementary Figure 7.**
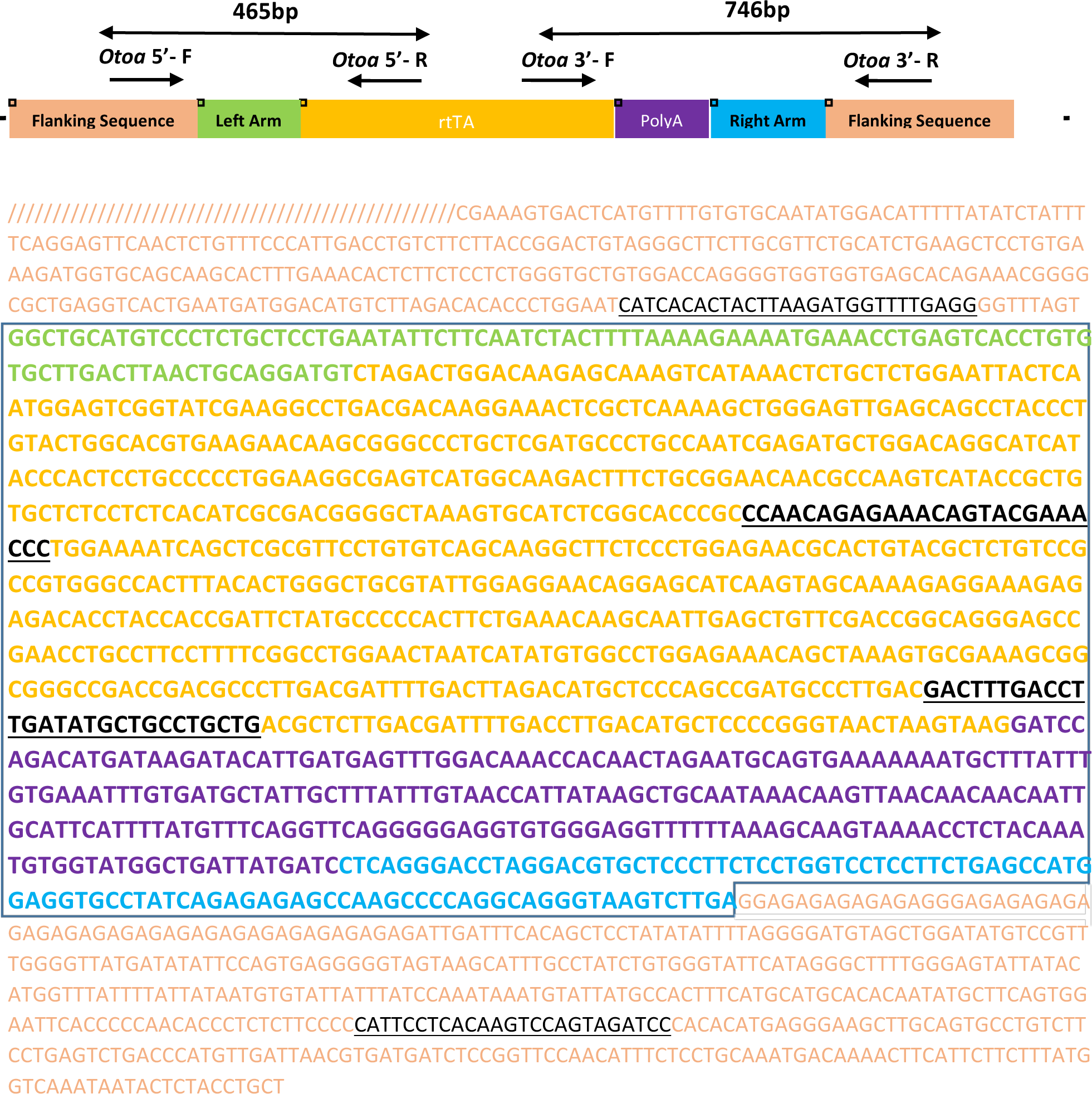
(a) Schematic and sequence of the *Otoa*-rtTA ssDNA. Various sequence elements are color coded. The primers used for 5′ and 3′ junction PCR and the amplicon sizes are shown above the schematic. The full sequence of the ssDNA cassette (including the homology arms) is boxed. Underlined black sequences are primer sequences.

**Supplementary Figure 8.**
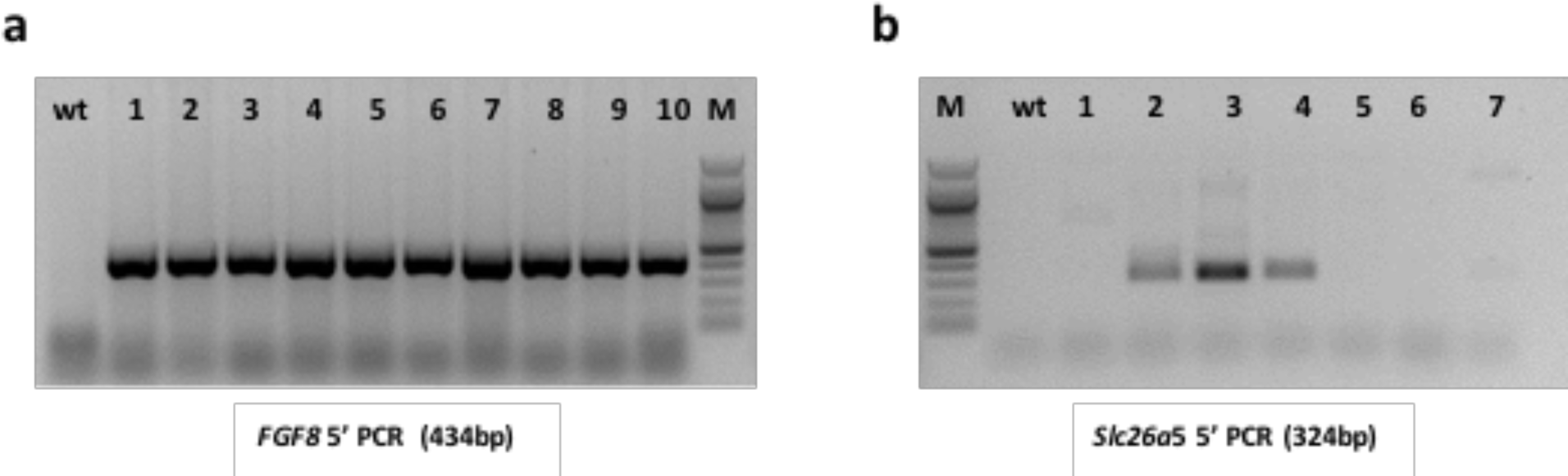
Germline transmission of *Easi*-CRISPR knock-in mouse models. **(a)** Genotyping of F1 pups derived from mating of the *Fgf8*-P2A-FlpO founder #4 (Figure 1f lane 4). Note that this founder is a homozygote and mating to a wild type mouse resulted in all pups heterozygous for the FlpO insertion. **(b)** Genotyping of F1 pups from *Slc25a5*-P2A-FlpO founder #3 (Supplementary Figure 3e lane 3) showing that F1 pups #2, #3 and #4 contain the knock-in cassette.

**Supplementary Figure 9.**
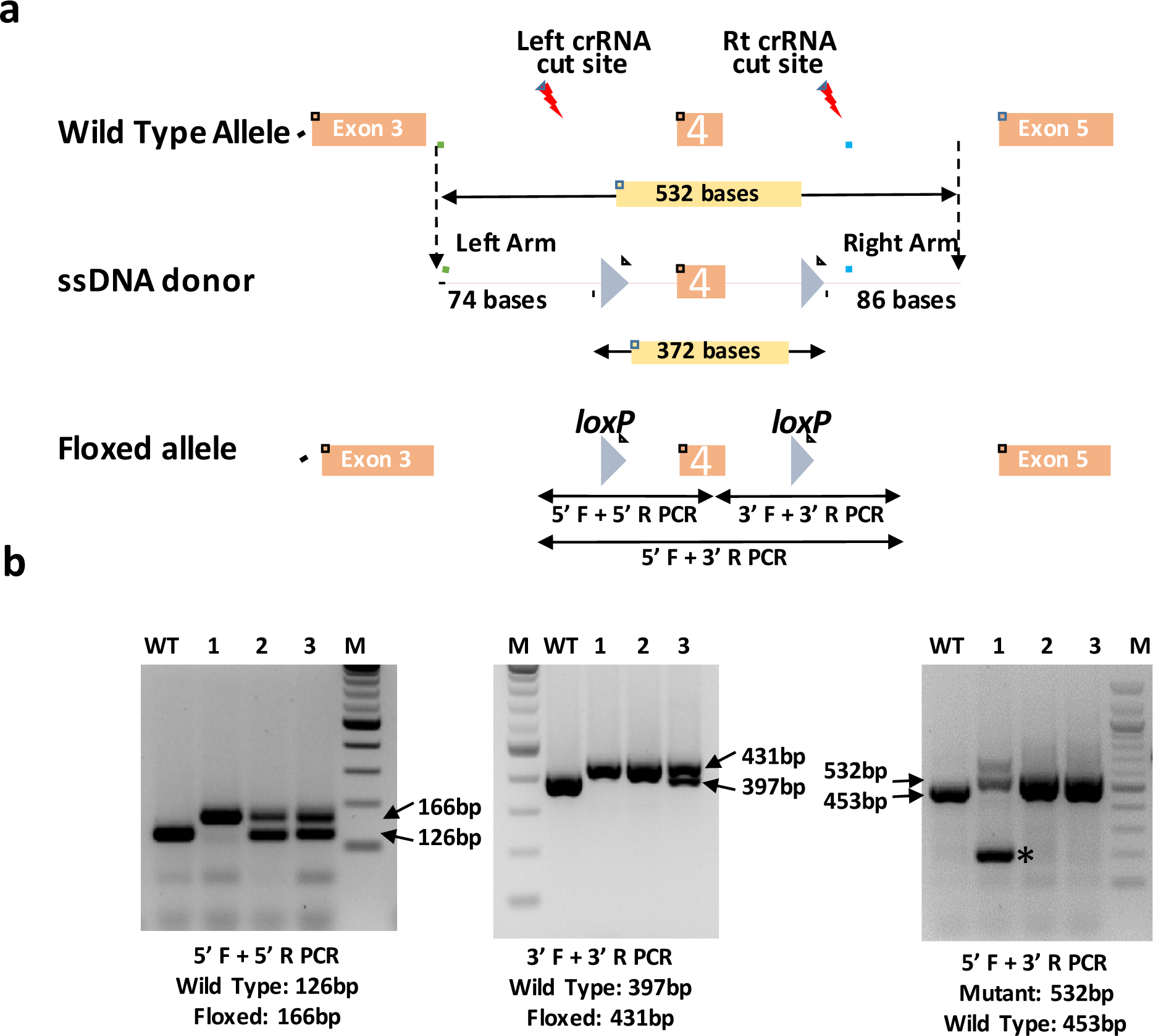
Generation of floxed *Paf1* allele using *Easi*-CRISPR. **(a)** Schematic showing wild type allele, ssDNA donor and targeted alleles. The lengths of ssDNA, homology arms and the distance between the two *loxP* sites are shown. **(b)** Genotyping of F0 animals. The expected sizes of PCR amplicons (wild type or floxed) are indicated. The gel on the right shows PCR bands amplified using the external primers to both *loxPs,* which shows that Pup #1 was heterozygous for both 5′ and 3′ *loxP* sites and the second allele had a deletion (shown by an asterisk); pup #2 was heterozygous for 5′ *loxP* and homozygous for 3′ *loxP* and the pup #3 was heterozygous for both *loxPs.*

**Supplementary Figure 10.**
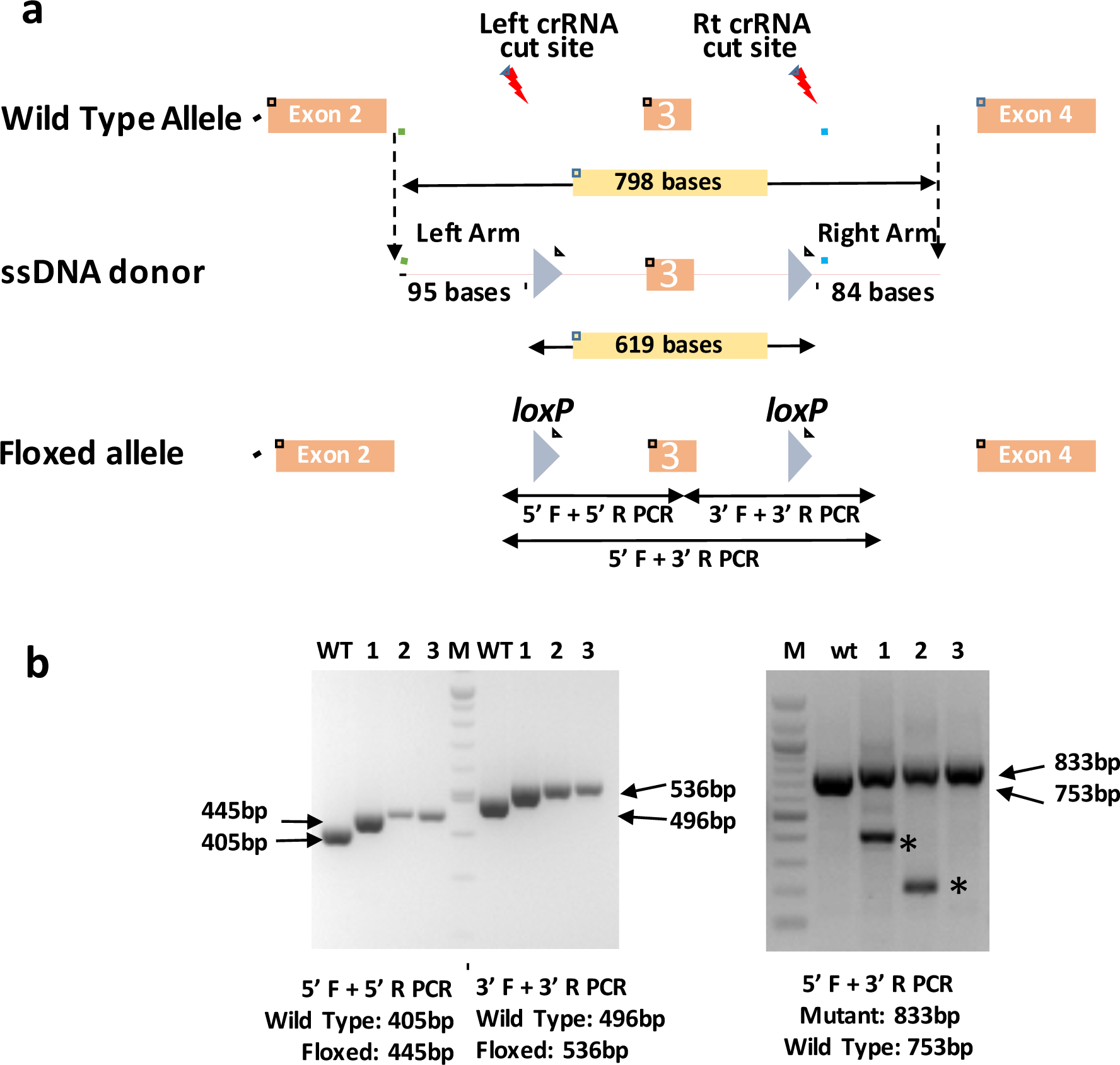
Generation of floxed *Ppp2r2a* allele using *Easi*-CRISPR. **(a)** Schematic showing wild type allele, ssDNA donor and targeted alleles. The lengths of ssDNA, homology arms and the distance between the two *loxP* sites are shown. **(b)** Genotyping of F0 animals for detection. The expected sizes of PCR amplicons (wild type or floxed) are indicated. The gel on the right shows PCR bands amplified using the external primers to both *loxPs* and Pup # 1 and 2 are heterozygous for both *loxPs* with deletions in their second alleles and the pup #3 is biallelic.

**Supplementary Figure 11.**
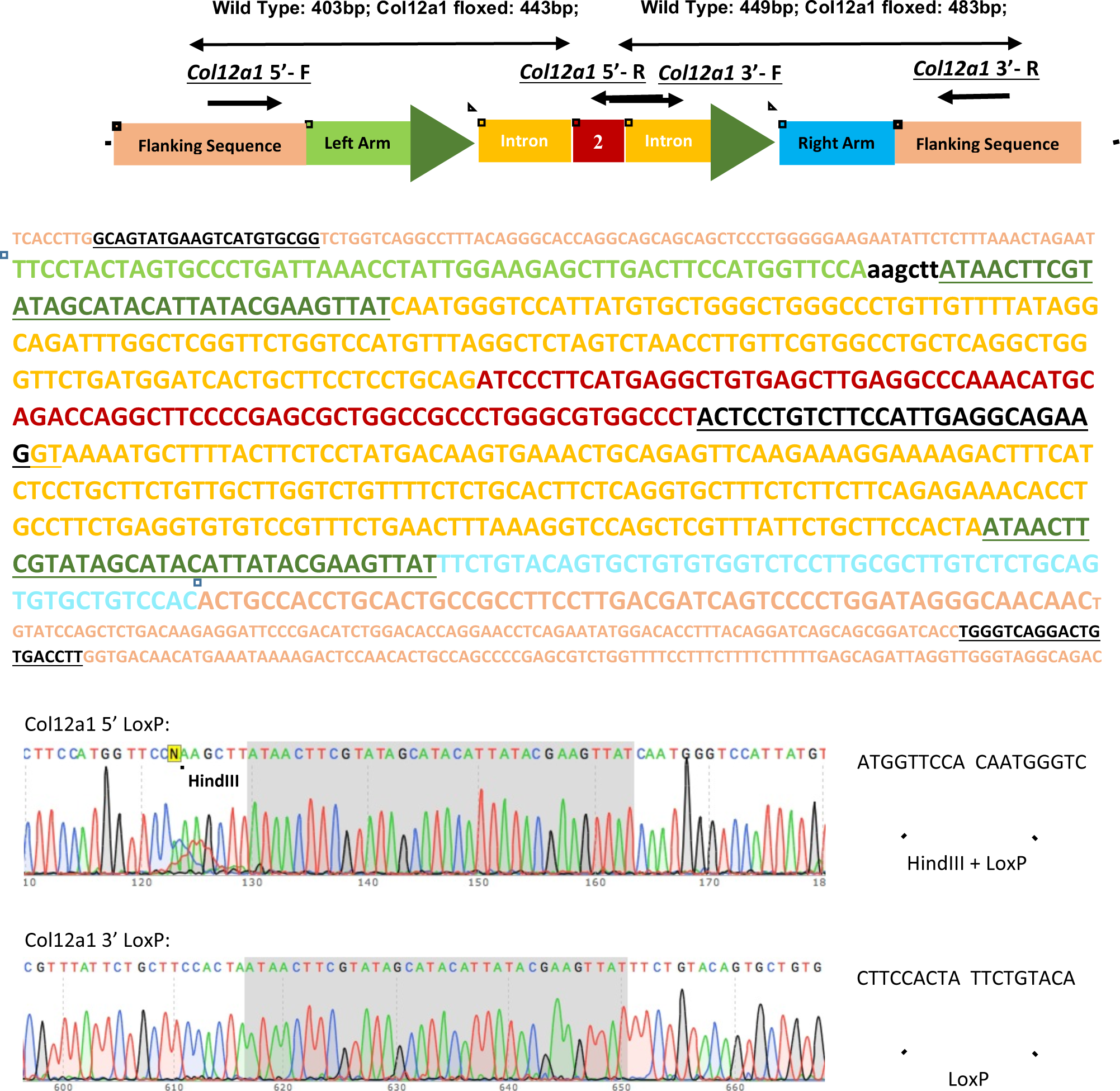
(a) Schematic and the sequence of *Col12a1* floxed ssDNA. Various sequence elements are color coded. The primers used for amplifying *loxP* insertion sites and their amplicon sizes are shown above the schematic. The full sequence of the ssDNA cassette (including the homology arms) is boxed. Underlined regions are primer sequences. (b) Sequencing of the *loxP* insertion sites a founder F0 pup showing correct insertion of the *loxP* sites and the HindIII site. The PCR amplicons were sequenced using one of the respective PCR primers.

**Supplementary Figure 12.**
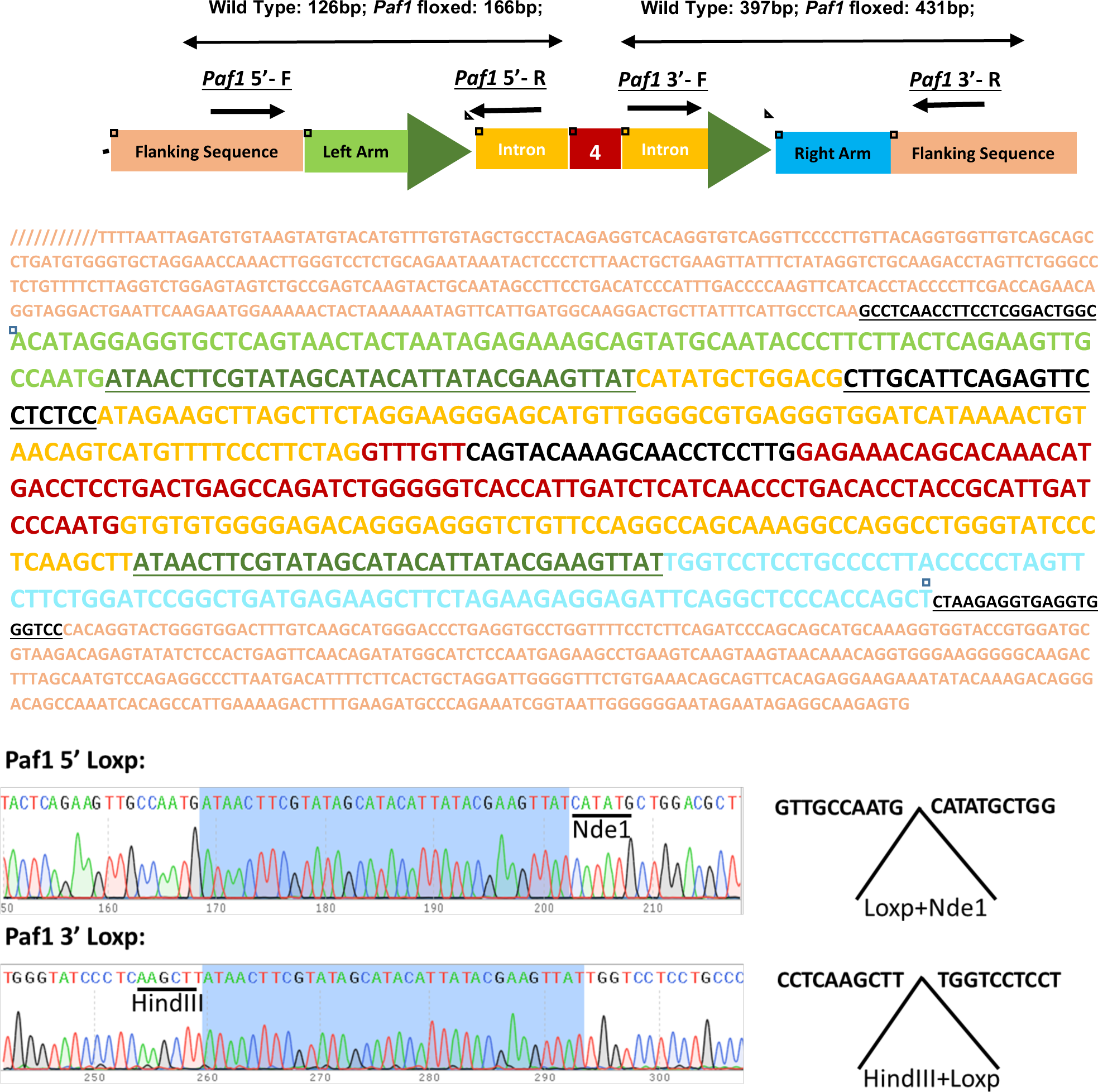
(a) Schematic and the sequence of *Paf-1* floxed ssDNA. Various sequence elements are color coded. The primers used for amplifying *loxP* insertion sites and their amplicon sizes are shown above the schematic. The full sequence of the ssDNA cassette (including the homology arms) is boxed. Underlined regions are primer sequences. (b) Sequencing of the *loxP* insertion sites a founder F0 pup showing correct insertion of the *loxP* sites and the NdeI and HindIII sites. The PCR amplicons were sequenced using one of the respective PCR primers.

**Supplementary Figure 13.**
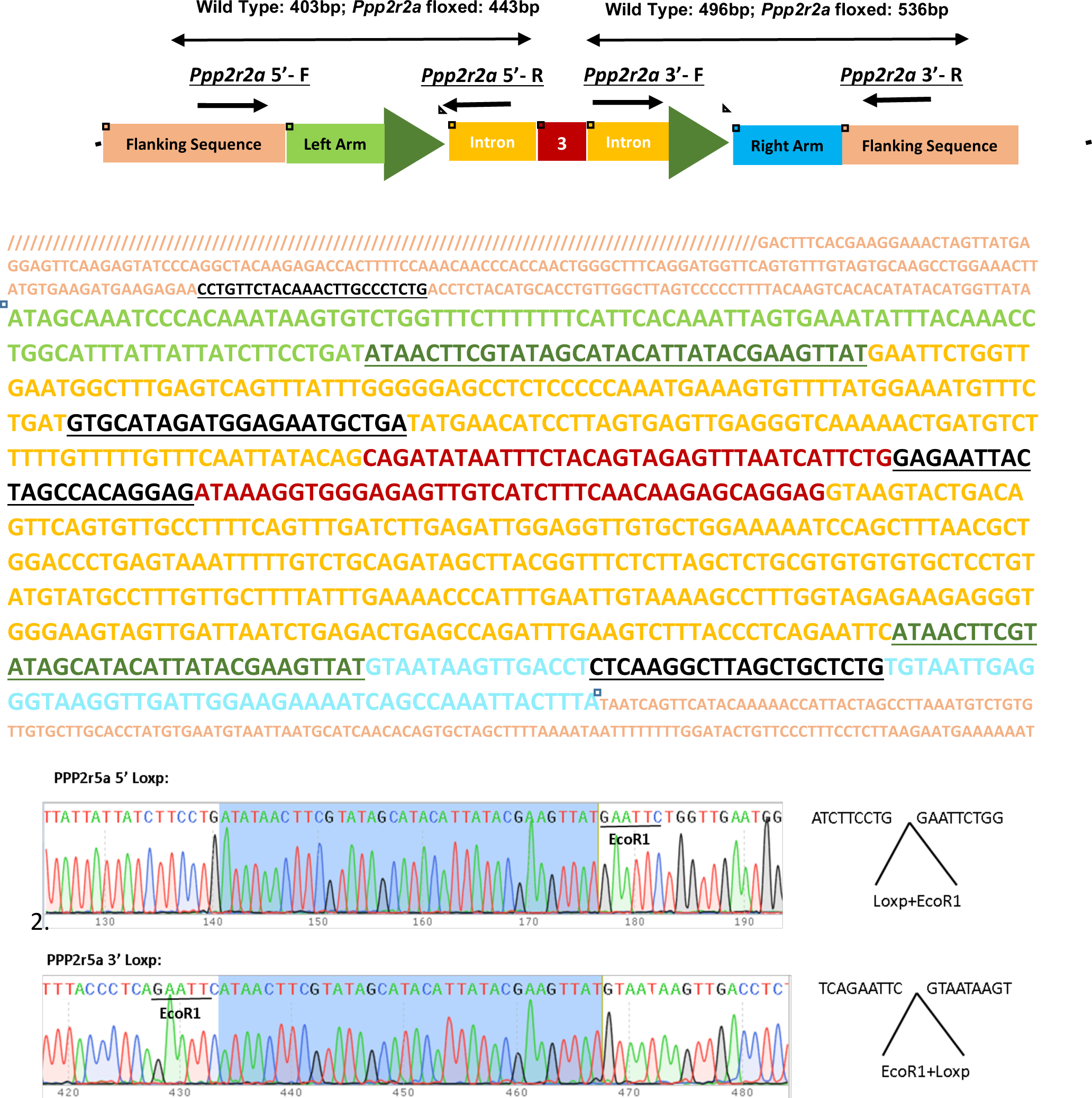
(a) Schematic and the sequence of *Ppp2r2a* floxed ssDNA. Various sequence elements are color coded. The primers used for amplifying *loxP* insertion sites and their amplicon sizes are shown above the schematic. The full sequence of the ssDNA cassette (including the homology arms) is boxed. Underlined regions are primer sequences. (b) Sequencing of the *loxP* insertion sites in a founder F0 pup showing correct insertion of the *loxP* sites and the EcoRI sites. The PCR amplicons were sequenced using one of the respective PCR primers.

